# A time irreversible model of nucleotide substitution for the SARS-CoV-2 evolution

**DOI:** 10.1101/2021.08.16.456444

**Authors:** Kazuharu Misawa

## Abstract

SARS-CoV-2 is the cause of the worldwide epidemic of severe acute respiratory syndrome. Evolutionary studies of the virus genome will provide a predictor of the fate of COVID-19 in the near future. Recent studies of the virus genomes have shown that C to U substitutions are overrepresented in the genome sequences of SARS-CoV-2. Traditional time-reversible substitution models cannot be applied to the evolution of SARS-CoV-2 sequences. Therefore, in this study, I propose a new time-irreversible model and a new method for estimating the nucleotide substitution rate of SARS-CoV-2. Computer simulations showed that that the new method gives good estimates. I applied the new method to estimate nucleotide substitution rates of SARS-CoV-2 sequences. The result suggests that the rate of C to U substitution of SARS-Cov-2 is ten times higher than other types of substitutions.

## 1 Introduction

Severe acute respiratory syndrome coronavirus-2 (SARS-CoV-2) is an RNA virus that has spready globally and is the cause of the COVID-19 pandemic. Evolutionary studies of the virus genome will provide a predictor of the fate of COVID-19 in the near future. Thus, it is important to estimate the nucleotide substitution rates of SARS-Cov-2.

Genomic analyses of SARS-CoV-2 have demonstrated that 50% of the sequence mutations are C to U transitions, with an 8-fold base-frequency normalized directional asymmetry between C to U and U to C substitutions (Simmonds 2020; Iwasaki, et al. 2021). Time reversible methods, such as Jukes-Cantor model (1969), Kimura 2-parameter model (Kimura 1980), Hasegawa-Kishino-Yano model (Hasegawa, et al. 1985), Tamura and Nei model (Tamura and Nei 1993), and General Time Reversible model (Tamura and Nei 1993), cannot be applied to study evolution of SARS-CoV-2, because C to U substitution of SARS-CoV-2 is not time-reversible.

In this study, I propose a new model of nucleotide substitutions that is applicable in cases where the C to U substitution rate is high. Using this new model, the evolutionary rate of SARS-CoV-2 was estimated. SARS-CoV-2 sequences were used to estimate the parameters of the evolutionary process.

## 2 Materials and Methods

### 2.1 Theoretical Background

The four bases C, U, G, and A of RNA are designated 1, 2, 3, and 4, respectively. *P(t)* is the probability matrix whose *ij*-th element is *p(t, i, j)*, where each individual entry, *p(t, i, j)* refers to the probability that nucleotide *i* is substituted by in time *t*. *N(t, i, j)* represents the numbers of the cases where the ancestral nucleotide is *i* and the derived nucleotide is *j* in time *t*. *N(t)* is the matrix whose *ij*-th element is *N(t, i, j)*.

The expected value of *N(t)* can be obtained by

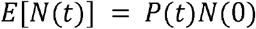

where *N*(0) is diag (*n*(1), *n*(2), *n*(3), *n*(4) and *n(i)* is the number of nucleotides *i* in the ancestral sequence.

Substitution rate matrix is a derivative of the substitution probability matrix with respect to time *t*.

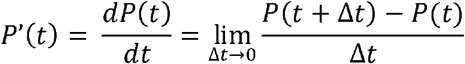

A rate matrix, R, is substituted by nucleotide *j* is defined by

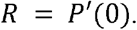

In this study, the process of nucleotide substitution is considered as a continuous Markov process.

### 2.2 Time Irreversible Model

To model the directional asymmetry between C to U and U to C substitutions, the following matrix *R* is proposed:

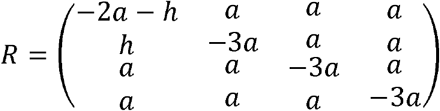

Let us define another matrix *Q* by

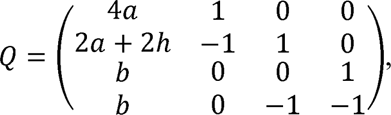

where *b* = 3*a* + *h*. It is worth noting that *R* can be diagonalized by *Q*

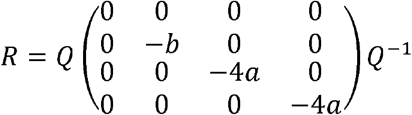

The matrix *P*(*t*) satisfies the Chapman-Kolmogorov equation:

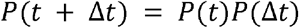

Thus, we obtain

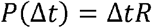

Therefore:

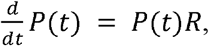

and

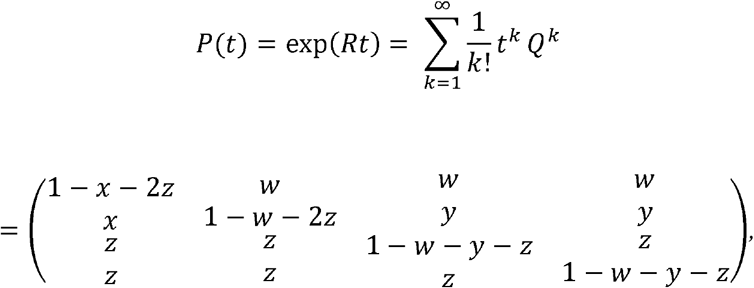

where

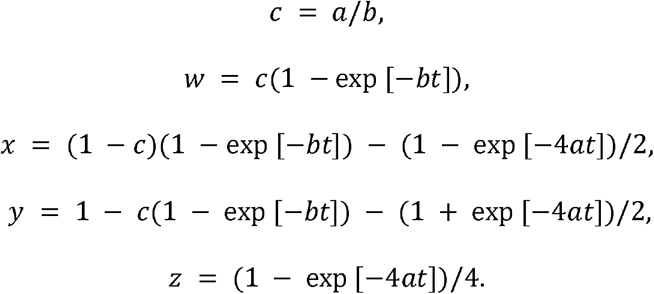

It is worth noting that there is an identity for all *t*.

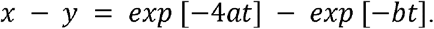

### 2.3 Estimatiion

The estimate of *P(t)*, 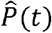, is obtained by

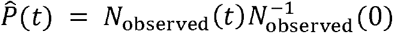

*w, x, y*, and *z* can be estimated from the elements of 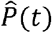. When multiple estimates are obtained, the arithmetic mean is used. By solve simultaneous equation *at* and *bt* can be estimated. Standard errors of these estimates can be obtained by the bootstrap method.

### 2.4 Simulation

Computer simulation was conducted by assuming the *a* = 0.001 and *h* = 10*a*. Transitions and transversions were estimated by Tamura and Nei (Tamura and Nei 1993) method using MEGA-X (Kumar, et al. 2018).

### 2.5 Sequence data

To estimate the substitution rate of nucleotides, a coronavirus genome found in South Africa was used (GISAID ID, EPI_ISL_736979). The sampling date was December 9, 2020. The Wuhan-Hu-1 reference genome sequence (GISAID ID, EPI_ISL_402125; GenBank ID, MN908947.3) sampled on December 31, 2019 (Wu, et al. 2020) was used as the ancestral virus. Sequences were aligned using MAFFT (Katoh, et al. 2002).

## 3 Results

Figure 1 shows the result of computer simulation. This figure shows that the new method gives good estimates.

**Figure 1.**
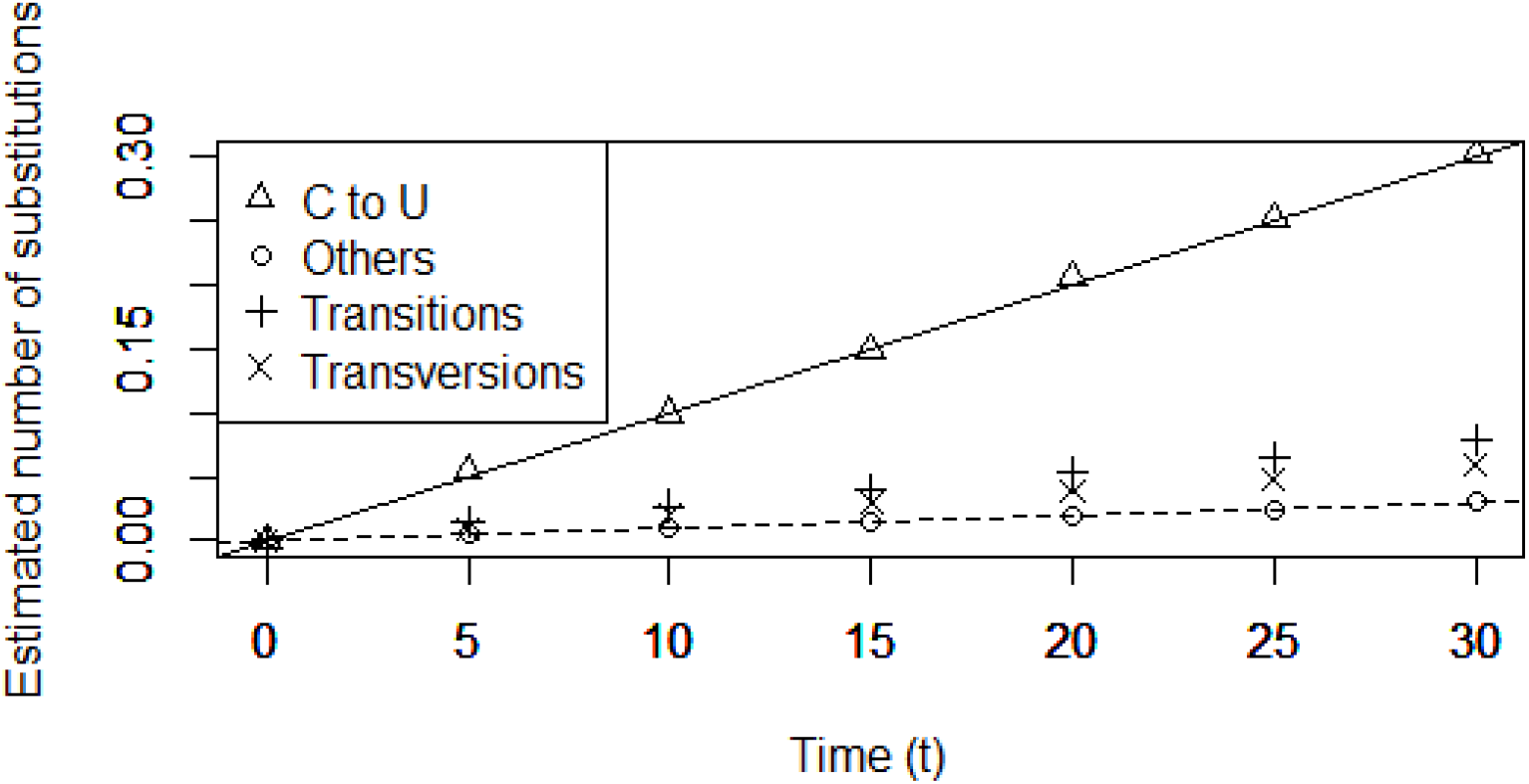
Computer simulation for estimating substitution rates The solid line shows the correct value of *ht* and the dashed line shows the correct value of *at*. Open triangle: The estimates of C to U substitutions obtained the new method. Open circle: the estimates of other types of substitutions obtained the new method. +: Estimated number of transitions obtained by Tamura-Nei method. ×: Estimated number of transversions obtained by Tamura-Nei method.

Table 1 shows the number of observed nucleotide differences Table 2 shows the estimates of the substitution rates. The rate of C to U substitution of SARS-Cov-2 is ten times higher than other types of substitution.

**Table 1.** Number of observed nucleotide differences.

| Descendant | Ancestor |  |  |  |
| --- | --- | --- | --- | --- |
|  | C | U | A | G |
| C | 5456 | 0 | 0 | 2 |
| U | 13 | 9528 | 4 | 2 |
| A | 0 | 0 | 5831 | 3 |
| G | 0 | 0 | 1 | 8907 |

**Table 2.** Estimates of substitution rates

| Type | Substitution rate | ± | Standard errors |
| --- | --- | --- | --- |
| C to U | $2.01 \times 10^{-3}$ | ± | $0.71 \times 10^{-3}$ |
| Others | $8.47 \times 10^{-5}$ | ± | $4.39 \times 10^{-5}$ |

## 4 Discussion

Because of high mutation rate from C to U, traditional time-reversible models cannot be used for SARS-CoV-2 evolution. To estimate the nucleotide substitution rates of SARS- Cov-2, a time-irreversible model is developed. Computer simulations showed that the new method gives good estimates even when the nucleotide substitution is not time-reversible.

Recent studies have shown that C to U substitutions of SARS-CoV-2 are overrepresented in the genome sequences of SARS-CoV-2. Table RNA molecules are subjected to cytosine-to-uracil editing through the action of members of the APOBEC family of cytidine deaminases (Bishop, et al. 2004). APOBEC-like directional C to U transitions of genomic plus-strand RNA are highly overrepresented in the SARS-CoV-2 genome sequences of variants that are emerging during the COVID-19 pandemic (Ratcliff and Simmonds 2021).

## Acknowledgment

I gratefully acknowledge institutes where genetic sequence data were generated and shared via the GISAID Initiative, on which this research is based. This work was supported by JSPS KAKENHI Grant Numbers JP19K22647,JP20K07316.

